# Functional analysis of a novel *de novo* variant in *PPP5C* associated with microcephaly, seizures, and developmental delay

**DOI:** 10.1101/2022.02.02.478908

**Authors:** Sara M. Fielder, Jill A. Rosenfeld, Lindsay C. Burrage, Lisa Emrick, Seema Lalani, Ruben Attali, Joshua N. Bembenek, Hieu Hoang, Dustin Baldridge, Gary A. Silverman, Undiagnosed Diseases Network, Tim Schedl, Stephen C. Pak

**Affiliations:** Department of Pediatrics, Washington University in St Louis School of Medicine, St Louis, MO 63110, USA; Department of Molecular and Human Genetics, Baylor College of Medicine, Houston, TX 77030, USA; Texas Children’s Hospital, Houston, TX 77030, USA; Genomic Research Department, Emedgene Technologies, 67443 Tel Aviv, Israel; Department of Obstetrics and Gynecology, C.S. Mott Center for Human Growth and Development, Wayne State University School of Medicine, Detroit, MI, 48201; Department of Genetics, Washington University in St Louis School of Medicine, St Louis, MO 63110, USA

**Keywords:** PPP5C, PPH-5, microcephaly, developmental delay, *C. elegans*

## Abstract

We describe a proband evaluated through the Undiagnosed Diseases Network (UDN) who presented with microcephaly, developmental delay, and refractory epilepsy with a *de novo* p.Ala47Thr missense variant in the protein phosphatase gene, *PPP5C*. This gene has not previously been associated with a Mendelian disease, and based on the population database, gnomAD, the gene has a low tolerance for loss-of-function variants (pLI=1, o/e=0.07). We functionally evaluated the *PPP5C* variant in *C. elegans* by knocking the variant into the orthologous gene, *pph-5,* at the corresponding residue, Ala48Thr. We employed assays in three different biological processes where *pph-5* was known to function through opposing the activity of genes, *mec-15* and *sep-1.* We demonstrated that, in contrast to control animals, the *pph-5* Ala48Thr variant suppresses the neurite growth phenotype and the GABA signaling defects of *mec-15* mutants, and the embryonic lethality of *sep-1* mutants. The Ala48Thr variant did not display dominance and behaved similarly to the reference *pph-5* null, indicating that the variant is likely a strong hypomorph or complete loss-of-function. We conclude that *pph-5* Ala48Thr is damaging in *C. elegans.* By extension in the proband, *PPP5C* p.Ala47Thr is likely damaging, the *de novo* dominant presentation is consistent with haplo-insufficiency, and the *PPP5C* variant is likely responsible for one or more of the proband’s phenotypes.

## INTRODUCTION

Protein phosphatase 5, encoded by the *PPP5C* gene, is a serine/threonine phosphatase that participates in the modulation of protein function by removing phosphate groups at serine and threonine residues. PPP5C interacts with a wide range of proteins, including nuclear receptors, kinases and transcription factors, and plays an important role in the regulation of numerous cellular processes including cell growth and differentiation, migration, apoptosis, DNA damage response and regulation of circadian rhythm [1–8] (reviewed in [9]). PPP5C is expressed ubiquitously but is present in high levels in the mammalian brain [10–13], and altered phosphatase activity has been associated with disease. For example, elevated PPP5C expression was detected in tumors, but not in normal tissues taken from patients with prostate cancer [14]. Additionally, knock down of PPP5C caused a decrease in the proliferation of cultured prostate, lung and urinary bladder cancer cells [5, 14, 15]. In contrast, overexpression of PPP5C in a breast cancer cell line (MCF7) enabled estrogen-dependent cells to grow independently of estrogen [16]. PPP5C has also been shown to regulate ciliogenesis by dephosphorylating DVL2, a key mediator of WNT signaling, in mammalian cells [17]. PPP5C is known to physically interact with and dephosphorylate HSP90 and glucocorticoid receptors suggesting a role in metabolism, stress response, immune function, skeletal growth, reproduction and cognition [18, 19]. HSP90 is a protein chaperone responsible for stabilizing many different proteins. For example, in conjunction with the TEL2-TTI1-TTI2 (TTT) complex, HSP90 stabilizes phosphatidylinositol 3-kinase related kinases (PIKKs), which in turn are responsible for regulating DNA damage responses, cell growth, and transcriptome surveillance (reviewed in [20, 21]).

*Ppp5c* homozygous knockout mice are viable and fertile, however, these mice have lower body weights as compared to their wild type littermates and do not gain weight when fed a high-fat diet [22–24]. In *Drosophila,* the *PPP5C* homolog (PpD3) is most highly expressed during embryonic development and has been implicated in centrosome separation during mitosis [25, 26]. Ppt1p, the yeast homolog of *PPP5C,* shows peak expression during early log cell growth, suggesting that Ppt1p may be involved in cell growth similar to what has been reported in human cell culture [27].

Analogous to the mammalian PPP5C, the *Caenorhabditis elegans* (*C. elegans)* ortholog PPH-5 is known to associate with the HSP90 and the glucocorticoid receptor complex [2, 28]. While *pph-5* null animals are superficially wild-type, genetic studies in *C. elegans* have revealed a role for *pph-5* in the regulation of GABA signaling and in neurite growth by opposing the activity of the F-box protein, MEC-15 [28]. Similarly, numerous *pph-5* loss-of-function mutants were identified in a genetic screen for interactors with separase (*sep-1*), a cysteine protease that plays an important role in chromosome segregation during mitosis, cortical granule exocytosis, and eggshell formation [29, 30]. In a *sep-1* hypomorphic mutant that is unable to form an eggshell, loss-of-function mutations in *pph-5* were found to rescue the embryonic lethality, suggesting a role for *pph-5* in the regulation of exocytosis and eggshell formation.

Here we present a proband accepted to the Undiagnosed Diseases Network (UDN) with a heterozygous, *de novo* missense variant (NM_006247.3, c.139G>A, p.Ala47Thr) in *PPP5C.* The proband’s main clinical features include microcephaly, epilepsy, and developmental delay. According to the population database, gnomAD v2.1.1 [31], *PPP5C* is highly constrained for loss-of-function variants (pLI=1.0, o/e=0.07). This gene has not previously been associated with a Mendelian disease. Therefore, we evaluated this gene-variant by performing *in vivo* functional analysis in the model organism *C. elegans*, as has been performed for other genes of uncertain significance [32–34]. We employed CRISPR-Cas9 to introduce the proband’s missense variant into *pph-5,* the *C. elegans* ortholog of *PPP5C*, and assessed the impact of *pph-5* Ala48Thr, corresponding to human *PPP5C* p.Ala47Thr. We provide data showing that the *pph-5* Ala48Thr variant alters neurite growth, GABA signaling, and eggshell formation, by acting as a strong hypomorph or loss-of-function allele. By extension, our data suggest that the human *PPP5C* p.Ala47Thr is damaging and likely responsible for one or more of the neurodevelopmental phenotypes in the proband.

## METHODS

### Proband evaluation and consent

Informed consent was obtained from the proband’s parents as part of enrollment into the Undiagnosed Diseases Network study, which is overseen by the National Institutes of Health Institutional Review Board (protocol #15HG0130). Phenotypic evaluation was completed through review of medical records and in-person examination by physicians who are co-investigators on the study.

### Exome sequencing and variant analysis

Clinical trio exome sequencing was performed clinically in a CLIA certified laboratory and the data was transferred for research analysis within the Undiagnosed Diseases Network. Codified Genomics (www.codifiedgenomics.com) and emedgene (www.emedgene.com) were used to prioritize novel, de novo variants that were not observed in gnomAD or in our database of over 1000 exomes and genomes (unless the phenotype of the probands matched) and biallelic variants in the coding region or near intron-exon boundaries with allele frequency of less than 1% and without homozygous entries in gnomAD [31] (Table S1). Variants of interest were confirmed using Sanger Sequencing within a CLIA certified laboratory.

### *C. elegans* strain and culture information

*C. elegans* were grown on standard Nematode Growth Media (NGM) and fed OP50 *E. coli* [35]. As *mec-15(u1042)* phenotypes were previously reported to be more extreme at higher temperatures [28], strains used for aldicarb assay and PLM-anterior neurite length were grown at 25°C for at least two generations before analysis. Strains used in this paper include (TU5237) *mec-15(u1042); uIs115* (a generous gift from the Chalfie lab), *pph-5(ok3498), sep-1(e2406)* and VC2010. Strains with *pph-5(udn90-94)* generated in this study are described below. A full list of strains used in this study is in Table S2. Some strains were provided by the CGC, which is funded by NIH Office of Research Infrastructure Programs (P40 OD010440).

### CRISPR/Cas9-based gene editing

*pph-5(udn90-94)* alleles were generated by injecting VC2010 1 day adults with Cas9 (IDT 1081060), tracrRNA (IDT 1072533), crRNA with the targeting sequence TGCAGATCAAGTGTACGACG (IDT Alt-R^®^ CRISPR-Cas9 sgRNA), pRF4 rol-6 plasmid as a positive injection marker, and single-stranded DNA oligonucleotide repair template (IDT) as previously described [32, 33]. Repair templates were 100 bases in length, centering on the locus of the proband variant, and were used to destroy the PAM site to prevent further editing, add a silent AvaII (NEB R0153L) restriction site to aid in genotyping, and to introduce the proband variant. Control repair templates containing only the silent PAM site change and the silent AvaII sites were also used, to determine that any phenotypes observed in the proband variant lines were due to the proband variant alone and not due to any silent mutations. Complete sequences of the repair templates are shown in Table S3. Each edited line was Sanger sequence verified and backcrossed twice to VC2010 to remove any background mutations. A line was considered independent if obtained from a separate injected animal. Three variant Ala48Thr lines were initially generated, and as all three lines showed no statistical difference in a preliminary blinded aldicarb paralysis assay (Fig. S1), only two lines were used for further analysis. Additionally, two control Ala48Ala edit lines were initially generated, and as both showed no statistical difference from each other or to the corresponding background strains in a blinded aldicarb paralysis assay and in the *sep-1* embryo lethality assay, only one control edit line was used for analysis.

### Aldicarb paralysis assay

To make 0.25mM aldicarb plates, 22.5 μL of 100mM aldicarb (Sigma #33386) stock in 70% ethanol was spread onto each NGM plate and allowed to dry at room temperature for one hour. A drop of OP50 suspended in LB was added to the center of each plate to help keep animals concentrated in the middle of the plate. Plates were then allowed to dry at room temperature for an additional hour. Ten one-day-old adult animals were added to each plate, with three replicates per strain. Paralyzed animals, defined as those not independently moving following two gentle taps with a platinum pick, were counted every twenty minutes. Animals were scored every thirty minutes from t=30 minutes to 130 minutes. Each experiment was performed three times, and the experimenter was blinded to genotypes. Plates that had two or more animals crawl off the agar were discarded from analysis.

### Microscopy and neurite length measurement

PLM <u>a</u>nterior <u>n</u>eurites (PLM-AN) tagged with RFP *(uls115[Pmec-17::RFP])* were imaged with a Zeiss compound fluorescence microscope. Strains were grown at 25°C for several generations before this assay was performed. Larval stage L4 animals were grown at 25°C for 24 hours, then immobilized in 2.5 mg/mL levamisole on dried 2% agarose pads. Length of the PLM-AN that had grown past the center of the vulva was measured (recorded as positive values), or PLM-AN that had not grown past the vulva, the length from the PLM-AN terminal to the center of the vulva was measured (recorded as negative values) using ImageJ. If both the bilaterally symmetric left and right PLM-ANs in a single animal were clearly visible and differentiable, both were measured. At least 37 PLM-AN lengths were recorded per line, from at least 23 animals.

### Obtaining heterozygote animals for phenotyping

To ensure heterozygous animals were generated, *mec-15; uIs115* larval stage L4 animals were raised on *fem-1* RNAi plates at 25°C, and females from the next generation were picked (i.e. 1 -day adults observed with stacked oocytes and no embryos that did not lay embryos for 4 hours). These females were mated to *pph-5(xxx); mec-15; uIs115* males, and resulting cross progeny were analyzed as described above. At least 29 PLMAN lengths were recorded per line, from at least 15 animals.

### Quantification of embryo lethality in *sep-1* mutant animals

Embryo lethality in *sep-1* separase and *pph-5* mutant animals were quantified as previously described [30]. In brief, strains were maintained at 15°C to obtain gravid mutant adult animals as *sep-1(e2406)* mutants have elevated sterility at higher temperatures. A single hermaphrodite animal staged between L4 and young adult was plated to individual 35mm NGM plates for 24 to 30 hours at 20°C and allowed to lay embryos. Animals were removed from the plate and embryos were incubated at 20°C for another 24 hours to allow hatching to occur. Dead embryos and live young larvae were counted to determine embryo lethality.

## RESULTS

### Clinical presentation

The proband was a 6-year-old girl with epilepsy, nystagmus, congenital microcephaly, and developmental delay (Fig. 1). Brain MRI initially at 11 months old showed slightly incomplete myelination that improved on an updated brain MRI at 4 years old but still with diminutive cerebral white matter volume (Fig. 1A-D). Her head size at birth was at the 2^nd^ percentile (32cm); by 18 months her head size was at −5.5 standard deviations (SD) and has remained in the −5.5SD to −6SD range (Fig. 1E). Seizure onset was at 12-16 months of age with generalized seizures including tonic clonic and drop seizures that are medically refractory. EEG has shown recurrent spike and slow-wave activity independently and multifocal spikes. Monitoring at age 4 showed subtle seizures every 2-13 minutes. Developmentally, she rolled over at 8-9 months with therapies, sat up after age 1 year, and currently crawls and pulls to a stand but is unable to walk independently. She is nonverbal and uses an augmentative communication device. On physical exam, she is microcephalic and has synophrys. Neurological exam noted bilateral horizontal nystagmus with inconsistent tracking but able to reach for objects. She is nonverbal, has decreased axial tone with increased tone in the extremities with scissoring in her legs with symmetric strength and increased deep tendon reflexes 3+ bilaterally in biceps, patellae and ankle clonus. She does not have any tremors, titubations or dysmetria on exam. She was also noted to have failure to thrive (Fig. 1F) and anemia. An ophthalmologic exam was significant for pendular nystagmus and moderate myopia. Family history is significant for a brother who passed away at birth; his pregnancy was complicated by pre-eclampsia, but the cause of death is unknown, and samples were unavailable for sequencing. Clinical trio exome sequencing was non-diagnostic. However, research analysis of the trio exome sequencing revealed a heterozygous, *de novo* missense variant in *PPP5C:* NM_006247.3:c. 139G>A: p.Ala47Thr. The variant has a CADD score of 29.4, is not found in gnomAD v2.1.1 [31], and the gene has not yet been associated with disease apart from a possible association with autism [36]. The gene level scores, pLI = 1 and o/e = 0.07, suggest a haploinsufficiency model for the genetic mechanism in humans.

**Figure 1.**
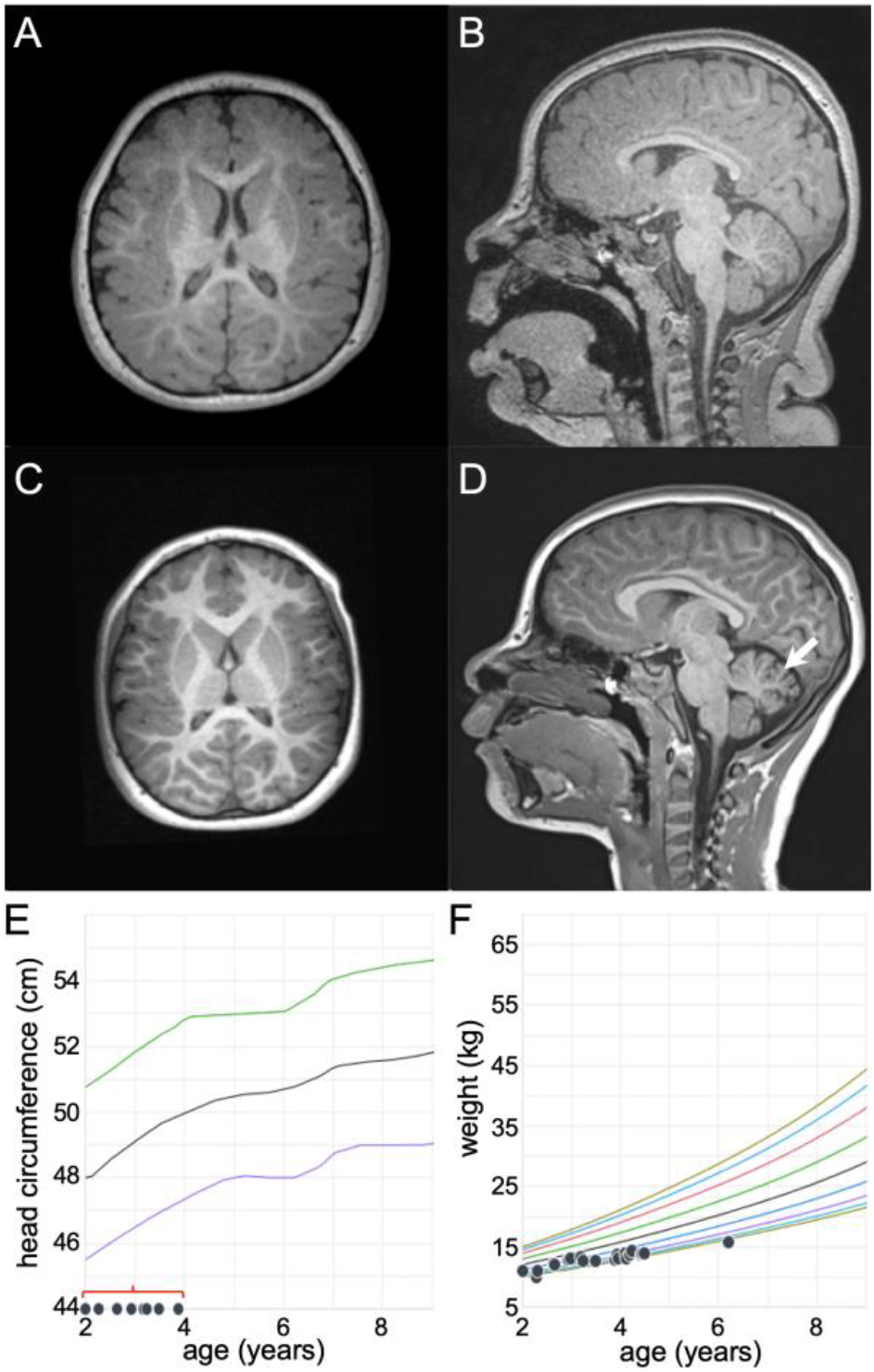
Proband brain MRI and growth chart. (A) T1 axial and (B) sagittal brain MRI at 11 months showing microcephaly with mild delay in myelination and decreased amount of white matter. (C) T1 axial and (D) sagittal brain MRI at 49 months demonstrating microcephaly with further myelination but persistent overall decreased amount of white matter. Additional findings on sagittal image showing decreased cerebellar folia indicative of mild cerebellar atrophy (white arrow). (E) Head circumference for age Nellhaus Girls 2 to 18 years old chart showing head circumference well below 2 standard deviations (red bracket) between 2 and 4 years of age. (F) Weight for age CDC chart demonstrating failure to thrive between 2 and 6 years old.

### *PPP5C* is conserved in *C. elegans*

To assess the effect of *PPP5C* p.Ala47Thr on *in vivo* function, we evaluated the impact of this gene variant in the model organism *C. elegans.* The ortholog of *PPP5C* in *C. elegans* is *pph-5,* which overall has an identity of 55.9% and a similarity of 72.3% to the human gene. In both organisms, *PPP5C* and *pph-5* encode two large domains: a phosphatase domain, which shares 62.7% identity (77.8% similarity), and three tetratricopeptide repeats (TPR) that are responsible for autoinhibition of the phosphatase domain by physically blocking the active site [37, 38]. The vast majority of mutations in *C. elegans pph-5* were previously identified in a suppressor screen [30] and are conserved or moderately conserved in humans (Table 1). The *PPP5C* p.Ala47Thr variant is located at the N-terminus of the protein in the TPR1 domain (Fig. 2A). The TPRs are highly conserved between *C. elegans* and humans, with PPH-5 and PPP5C sharing ~54% identity (72.5% similarity) in this domain (Fig. 2B). We therefore modeled the *PPP5C* p.Ala47Thr proband variant in the corresponding *C. elegans* residue (Ala48) of *pph-5* using CRISPR/Cas9-based gene editing. In addition to the variant residue, we introduced synonymous changes to block Cas9 re-cleavage following homology-directed repair and a restriction enzyme cleavage site to aid in allele genotyping. As controls, we generated lines that contain only the synonymous changes, but not the proband’s variant. In total, we generated three independent variant (Ala48Thr) alleles and two independent control (Ala48Ala) alleles (Table S2). All the CRISPR-edited lines were superficially wild-type, similar to the *pph-5(ok3498)* null mutants (hereafter referred to as *pph-5(null))* [29].

**Figure 2.**
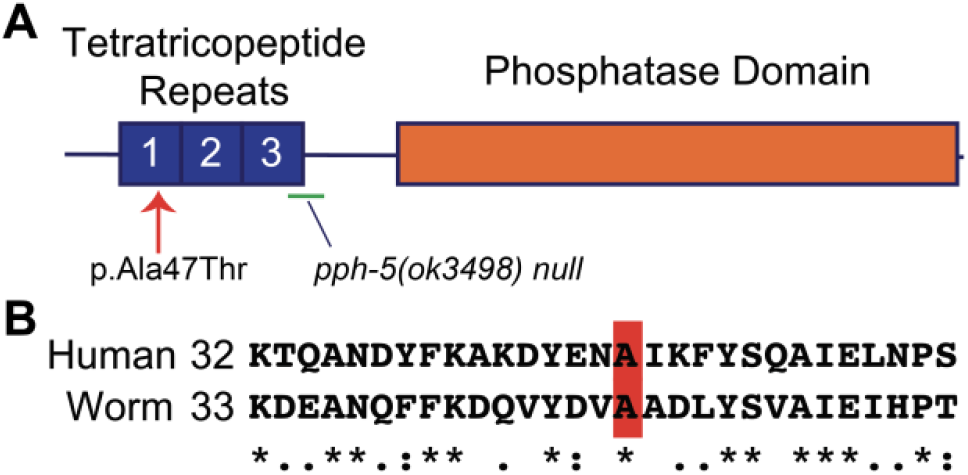
*pph-5* is the *C. elegans* ortholog of *PPP5C*. (A) PPP5C domain map, with the locations of the proband variant Ala47Thr (red arrow) and *C. elegans pph-5(null)* allele (green line) indicated. (B) Alignment of *H. sapiens* and *C. elegans* Tetratricopeptide repeat 1 region. Proband variant residue location highlighted in *red.*

**Table 1.** *pph-5* legacy missense alleles are experimental tests of yet to be observed *PPP5C* variants.

| Allele<sup>†</sup> | *pph-5*<br>residue | *pph-5*<br>change | Hypomorphic<br>character<sup>§</sup> | *PPP5C*<br>residue | gnomAD<br>instances<br>(n) | comments |
| --- | --- | --- | --- | --- | --- | --- |
| **Mutations in the TRP domain** |  |  |  |  |  |  |
| ***erb58*** | **Ile32** | **Asn** | **strong** | **Leu31** | **none** |  |
| ***erb47*** | **Tyr52** | **His** | **strong** | **Tyr51** | **none** |  |
| ***erb51*** | **Gly66** | **Glu** | **strong** | **Gly66** | **none** |  |
| *erb44* | Leu77 | Pro | weak | Cys77 | none | non-conservative change in residue |
| *erb54* | A105F | Phe | very weak | Ser105 | none | conservative change in residue |
| **Mutations in the phosphatase domain** |  |  |  |  |  |  |
| ***erb30*** | **Met211** | **Lys** | **strong** | **Ile212** | **Ile>Leu**<br>**(1)** | conservative change in residue. gnomAD<br>residue a conservative substitution |
| *erb1* | T229L | Leu | weak | T230 | none | conservative change in residue |
| ***erb46*** | **His243** | **Arg** | **strong** | **His244** | **none** |  |
| ***erb45*** | **Gly244** | **Glu** | **strong** | **Gly245** | **none** |  |
| ***erb31*** | **Asp270** | **Ala** | **strong** | **Asp271** | **none** |  |
| ***erb57*** | **Asp270** | **Asn** | **strong** | **Asp271** | **none** |  |
| *erb48* | Met285 | Arg | weak | Ile286 | Leu>Phe<br>(1) | conservative change in residue. gnomAD<br>residue a conservative substitution |
| *erb63* | Arg300 | Cys | weak | Arg301 | none |  |
| *erb21* | Asn309 | Lys | weak | Asn310 | Asn>Lys<br>(1) | presence in gnomAD may be explained by<br>the substitution being a weak hypomorph |
| *erb59* | Met311 | Arg | weak | Ile312 | none | conservative change in residue |
| *erb42* | Tyr322 | Ile | weak | Tyr323 | none |  |
| *erb72* | His351 | Arg | very weak | His352 | none |  |
| *av101* | Pro375 | Gln | very weak | Pro376 | none |  |
| *erb2* | Met380 | Thr | weak | Met381 | none |  |
| *erb68* | Ser397 | Pro | weak | Ser398 | none |  |
| *erb33* | Trp413 | Gly | weak | Phe414 | none | conservative change in residue |
| *erb38* | Cys414 | Tyr | weak | Leu415 | none | non-conservative change in residue |
| ***erb28*** | **His426** | **Gln** | **strong** | **His427** | **none** |  |
| ***erb22*** | **Cys441** | **Tyr** | **strong** | **Cys442** | **none** |  |
| *erb65* | Thr443 | Ile | weak | Thr444 | none |  |
| *erb40* | Pro448 | Leu | weak | Pro449 | none |  |
| *erb52* | Gly458 | Glu | weak | Ala459 | none | conservative change in residue |
| ***erb13*** | **Tyr65\*** | **stop** | **strong/null** | **Tyr64** | **none** |  |

<sup>†</sup> Mutations identified in *Richie et al., 2011* or *Melesse et al., 2018*. <sup>§</sup> Hypomorphic character accessed from suppression of *sep-1(e2406)* embryonic lethality. Bold indicates alleles that are strong suppressors of *sep-1(e2406)* embryonic lethality phenotype.

### *pph-5* Ala48Thr variant is defective in function and suppresses the touch receptor neuron neurite growth phenotype in *mec-15* F-box protein mutants

Since the *pph-5* Ala48Thr proband variant and the *pph-5* null mutants were superficially wild-type, we employed sensitized genetic backgrounds, the *mec-15* null or the *sep-1* hypomorph, where it is known that *pph-5* acts to oppose the function of these genes. In *C. elegans, pph-5* has previously been reported to have genetic interactions with *mec-15* in regulating neurite growth [28]. MEC-15 is an F-box protein (orthologous to human *FBXW9)* that promotes touch receptor neurite growth by aiding degradation of DLK-1 (orthologous to human *MAP3K12),* an inhibitor of microtubule stability [28, 39]. As a consequence, in *mec-15(u1042)* mutants, the microtubules are less stable, and the PLM anterior neurites (PLM-AN) are considerably shorter than those of wild-type animals. PPH-5 is part of the HSP90 chaperone complex, which aids in protein folding [40, 41], including that of DLK-1 [28]. Thus, PPH-5 negatively regulates neurite growth by opposing the function of MEC-15. Loss-of-function alleles of *pph-5,* and other components of the HSP90 complex, therefore restore microtubule stability in *mec-15(u1042)* mutants, and suppress the short PLM-AN phenotype [28]. To examine the role of the PPH-5 Ala48Thr proband variant in neurite growth, we crossed the *pph-5* variants into the *mec-15(u1042)* mutant background and measured PLM-AN neurite length. As previously reported, *mec-15* mutants have significantly shorter PLM-AN neurites compared to wild-type animals (Fig. 3A and B). However, when combined with the proband variant, the short neurite phenotype was completely suppressed and neurite length was restored to that of the wildtype (Fig. 3A and B). In contrast, the *pph-5* Ala48Ala control edited lines did not suppress the short neurite phenotype of *mec-15(u1042)* animals. These results indicate that *pph-5* Ala48Thr is defective in function as a negative regulator of PLM-AN neurite length. Moreover, the extent of the suppression by the proband’s variant was similar to that of the reference null allele of *pph-5,* indicating that the *pph-5* Ala48Thr variant is likely a strong hypomorph or a complete loss-of-function.

**Figure 3.**
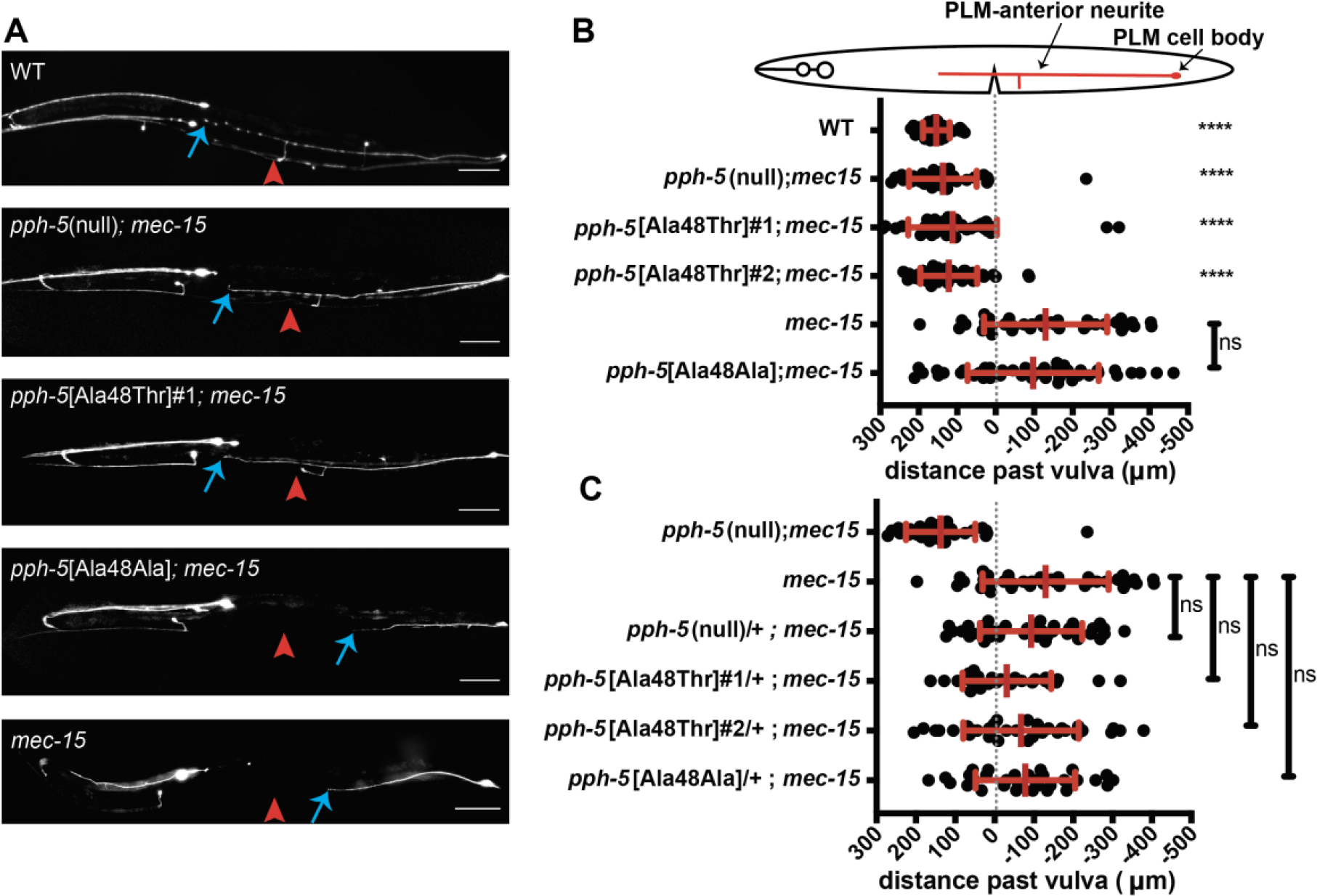
Ala48Thr variant suppresses *mec-15* PLM-anterior neurite growth defect. (A) Images of day 1 adult animals expressing *pmec-17::RFP* in the PLM neurons. The PLM-anterior neurite normally grows (PLMAN terminus blue arrow) past the vulva (red arrowhead). However, *mec-15(u1042)* animals have a PLM-AN that typically does not grow past the vulva. Two independently generated lines of *pph-5[Ala48Thr]; mec-15* animals have neurites that grow past the vulva, while the control *pph-5[Ala48Ala]; mec-15* mimics the *mec-15* PLM-AN length. (B) Quantification of neurite termination in relation to the vulva of at least 37 neurites from at least 23 animals per genotype. Negative values indicate PLM-AN did not grow past the vulva, and indicate distance from PLM-AN terminus to vulva, while positive values indicate length the PLM-AN grew past the vulva in the anterior direction. Vulval location indicated by dashed vertical line. Mean and standard deviation plotted. Statistical significance indicated is genotype as compared to *mec-15.* (C) Quantification of at least 29 neurites from at least 15 heterozygous *pph-5;* homozygous *mec-15(u1042)* animals. Kruskall-Wallis ANOVA was performed, followed by post-hoc Dunn’s multiple comparisons tests in both B and C. (ns, **** p ≤ 0.0001).

### *pph-5* Ala48Thr variant suppresses aldicarb-induced paralysis of *mec-15* mutants

In the *C. elegans* neuromuscular junction, opposing neurotransmitter signals regulate muscle contraction. Acetylcholine (ACh) provides the stimulatory signal for muscle contraction while gamma-aminobutyric acid (GABA) provides the inhibitory signal. ACh activity is regulated by acetylcholinesterase, an enzyme that degrades ACh in the neuromuscular junction. Inhibition of acetylcholinesterase by treatment with the drug aldicarb leads to ACh accumulation in the extracellular space and subsequent hypercontraction of the muscle and eventual paralysis (reviewed in [42]). MEC-15 promotes synaptic transmission in GABAergic motor neurons to inhibit muscle contraction [43, 44]. Consequently, when exposed to aldicarb, *mec-15(u1042)* mutants release less GABA and reach paralysis at a faster rate than wild-type animals (Fig. 4A) [28, 44, 45]. This increased aldicarb sensitivity of *mec-15* mutants can be suppressed by loss-of-function mutations in *pph-5,* indicating that PPH-5 negatively regulates the release of GABA [28]. Thus, we used an aldicarb sensitivity assay to assess the impact of *pph-5* Ala48Thr on GABA signaling. Upon aldicarb treatment, the time taken for 50% of *mec-15(u1042)* animals to reach paralysis was 62 minutes, which was similar to that of the control edit *pph-5*[Ala48Ala]*; mec-15(u1042)* animals (Fig. 4B). In contrast, the *pph-5*[Ala48Thr]*; mec-15(u1042)* animals were less sensitive to aldicarb, and took 86 minutes to reach 50% paralysis. These results were indistinguishable to those obtained with the *pph-5(null); mec-15(u1042)* animals (Fig. 4B). Taken together, our aldicarb-sensitivity data provide additional evidence for *pph-5* Ala48Thr variant being damaging to gene function and behaving similarly to the null allele.

**Figure 4.**
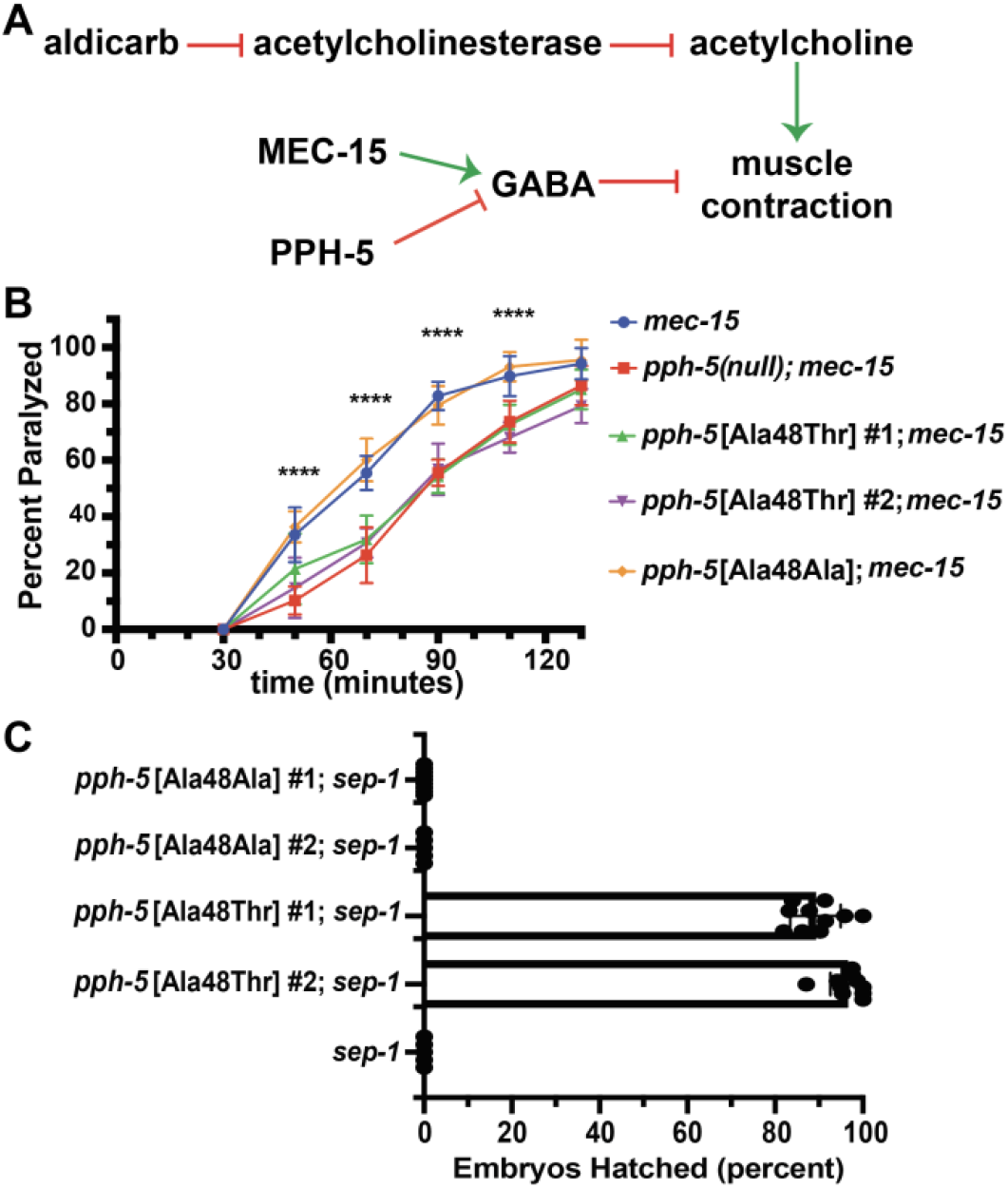
Ala48Thr variant suppresses known *pph-5* interaction phenotypes. (A) A genetic model of PPH-5/MEC-15 interaction in GABA signaling, (B) Quantification of percent of animals paralyzed at the indicated time after the start of exposure to aldicarb. Statistical significance as compared to *mec-15.* t-tests (ns, **** p ≤ 0.0001). (C) Percent of embryos that hatched when *pph-5*[Ala48Thr] or *pph-5*[Ala48Ala] was combined with *sep-1(e2406).*

### *pph-5* Ala48Thr variant suppresses embryonic lethality of *sep-1* mutants

As a final test of the proband’s variant function, we examined the role of PPH-5 Ala48Thr in embryogenesis. *C. elegans sep-1* (orthologous to human *ESPL1)* encodes a cysteine protease required for the cleavage of cohesin complex subunits during mitosis, exocytosis of cortical granules and eggshell formation [46, 47]. Consequently, *sep-1(e2406)* hypomorphic mutants do not form an eggshell, fail to hatch, and are embryonic lethal [47]. However, reduction of function mutations in *pph-5,* including mutations within the TPR domain, are known to suppress the embryonic lethality of *sep-1* mutants, indicating that PPH-5 phosphatase activity has a role in negatively regulating both SEP-1 and eggshell formation [30]. Therefore, we used the suppression of embryonic lethality as a readout to assess the functionality of the proband’s variant. Consistent with previous results, we found *sep-1* animals to be 100% embryonic lethal (Fig. 4C). When the control edit *pph-5* Ala48Ala allele was combined with the *sep-1* allele, the resulting animals were also 100% embryonic lethal as indicated by the lack of hatched embryos (Fig. 4C). In contrast, embryonic lethality was strongly suppressed by the *pph-5* Ala48Thr variant and 89-96% of the embryos successfully hatched (Fig. 4C). The percent hatched values are similar to the previous data obtained for the *pph-5(tm2979)* null; *sep-1(e2406)* double mutant [30]. Combined, our results indicate that Ala48Thr variant leads to defective PPH-5 function with a severity that is similar to that of the *pph-5* null alleles.

### *pph-5* null and Ala48Thr variants behave recessively in *C. elegans*

The underrepresentation of loss-of-function variants in the control human population (gnomAD v2.1.1, pLI=1, o/e = 0.07) suggests that *PPP5C* is a haploinsufficient gene. However, missense variants in haploinsufficient genes may act in a gain-of-function manner, for example, via a dominant negative or a hyperactive mechanism. We therefore used the PLM-AN neurite growth assay to compare heterozygous and homozygous phenotypes for the *pph-5* null and Ala48Thr alleles.

In the *mec-15(u1042)* mutant background, the *pph-5 null* and *pph-5* Ala48Thr homozygous animals have normal PLM-AN neurite lengths (Fig. 3B), similar to wild-type animals indicating full suppression of *mec-15* phenotype. In contrast, animals heterozygous for *pph-5* Ala48Thr failed to suppress the *mec-15* mutation and had similar short neurite lengths compared to the *pph-5* Ala48Ala control edited animals (p=0.31) (Fig. 3C), suggesting that PPH-5 Ala48Thr does not have dominant gain-of-function activity. Similarly, animals heterozygous for *pph-5* null also fail to suppress the *mec-15* mutation, consistent with *pph-5* not being a haploinsufficient gene in *C. elegans*. The absence of haploinsufficiency for *pph-5*, compared to *PPP5C*, is not unexpected given that different species titrate gene expression differently and haploinsufficiency is thought to be rare in *C. elegans* (Hodgkin, 2005).

### *pph-5* legacy mutations as functional tests of yet to be identified *PPP5C* missense variants

A collection of 25 *pph-5* missense mutations has been generated in *C. elegans* forward genetic screens [29, 30]. The *pph-5* mutations were isolated following chemical mutagenesis as recessive suppressors of the *sep-1* separase embryonic lethality phenotype, analogous to the assay shown in Figure 4C. These *pph-5* legacy mutations may have already been observed or have yet to be observed in the human ortholog and provide pre-existing experimental functional information about conserved residues and variants in *PPP5C*.

We mapped the 25 legacy missense variants to the corresponding amino acid residues in *PPP5C* (Table 1). All the mutations reside in the TRP repeats or phosphatase domain. The wild type residue of the *pph-5* mutation is identical in *PPP5C* in 18 cases, while for the remaining seven, five contain conservative substitutions. From examination of the gnomAD population database, there are no examples of variants observed for 22 of the *PPP5C* residues, indicating that these residues show some level of constraint to variation. For two of the exceptions, PPP5C has conservative changes, and the variant residues are conservative substitutions. From the mutant screens, it was found that a modest reduction in PPH-5 function can suppress some *sep-1* mutant phenotypes, indicating that the screen is very sensitive [30]. We therefore subdivided the *pph-5* mutations into three categories based on the extent of suppression of *sep-1* mutant embryonic lethality [29, 30]: strong hypomorphs that are similar to null alleles (greater than 70% suppression); weak hypomorphs (between 30 – 70% suppression); and very weak hypomorphs (less than 30% suppression). Assuming haploinsufficiney for the *PPP5C* p.Ala47Thr dominant proband phenotypes, it is likely that the *pph-5* weak and very weak mutations do not reduce gene function enough to achieve haploinsufficiency in *PPP5C.* Therefore, we only considered the strong hypomorph *pph-5* mutations as sufficiently damaging to be similar to *pph-5* Ala48Thr. This analysis yielded 10 legacy *pph-5* strong hypomorphic mutations, where the residue is identical or a conservative substitution in *PPP5C*, and there are no variants in gnomAD (Table 1). The *C. elegans* functional analysis indicates that these 10 variants are damaging and, by extension, if observed as *PPP5C* heterozygotes are likely to impact human phenotype.

## DISCUSSION

In a pediatric proband with congenital microcephaly, epilepsy, and developmental delays, we identified a *de novo* heterozygous missense variant p.Ala47Thr in *PPP5C*, a gene not previously associated with Mendelian disease. By modeling the variant in the *C. elegans* ortholog, *pph-5,* we demonstrated that *pph-5* Ala48Thr is damaging to function *in vivo* using three different biological assays. In addition, the proband variant behaved similarly to the *pph-5* null allele indicating that *pph-5* Ala48Thr, and by extension *PPP5C* p.Ala47Thr, likely acts as a strong hypomorph or complete loss-of-function. These functional data, combined with the *in silico* prediction of likely deleterious (CADD score 29.4), the intolerance of *PPP5C* to loss of function variants (pL1=1, o/e=0.07), the absence of variants at residue p.Ala48 in control population databases (gnomAD v2.1.1), and the dominant presentation of the proband are consistent with a haploinsufficient mechanism for the proband’s phenotypes.

Although *pph-5* null and Ala48Thr variant animals are superficially wild-type, the variant function could be assessed in *C. elegans* using sensitized genetic backgrounds. MEC-15 promotes PLM-AN neurite growth by regulating the degradation of a HSP90-client protein, DLK-1, and thus, *mec-15* mutants have short neurites. Using the short PLM-AN neurite length of *mec-15* mutants as our readout, we determined that *pph-5* Ala48Thr is defective in its ability to negatively regulate touch receptor neurite growth (Fig. 3). Using similar suppression assays, we showed that *pph-5* Ala48Thr suppresses the aldicarb hypersensitivity of *mec-15* mutants (Fig. 4), indicating that the *pph-5* variant is defective in inhibition of GABA signaling. These *C. elegans* nervous system functions of *pph-5* are consistent with the neurodevelopmental phenotypes observed in the proband.

A recent large-scale exome sequencing study of 11,986 individuals with autism spectrum disorder (ASD) identified *PPP5C* as one of 102 risk genes [36]. These three reported probands with autism have variants of uncertain significance in the C-terminus of PPP5C, as compared to the proband described here with a variant in the N-terminal TPR domain. However, functional data evaluating the role of *PPP5C* missense variants in ASD and/or other neurodevelopmental disorders is currently lacking. *In vivo* gene function studies in model organisms like *C. elegans* offer a fast, cost-effective and quantitative approach in assessing the biological effects of future *de novo* variants identified in *PPP5C.*

The Ala47Thr variant is located in the TPR domain of PPP5C, which is thought to play an important role in the autoinhibition of the phosphatase activity through intramolecular interactions with the C-terminal domain that hinder substrate access to the phosphatase catalytic pocket [41]. The binding of HSP90 to the TPR domain of PPP5C relieves this autoinhibition and stimulates phosphatase activity. The Ala47Thr variant might disrupt binding with HSP90. The active phosphatase can subsequently activate one of several hundred known HSP90 client proteins [48]. One such client is ataxia and telangiectasia and Rad3 related (ATR), a kinase that activates DNA damage signaling response [48, 49]. Mutations in *ATR*, and other ATR-mediated checkpoint response genes, *MCPH1* and *ATRIP,* have been linked to microcephaly and specifically Seckel syndrome [50] (reviewed in [51]). This is thought to be because ATR has an important role in supporting the rapidly proliferating zones in the brains of mice [52, 53]. ATR, as well as other PIKK genes, are stabilized by the Hsp90 complex, through interaction with the Tel2-Tti1-Tti2 (TTT) complex [49, 54, 55]. Compound heterozygous variants in *TELO2* (the human homolog of yeast Tel2) have been identified in six probands, including three siblings and three unrelated individuals, with You-Hoover-Fong syndrome, which is characterized by microcephaly, movement disorder, and severely delayed development [56]. Recently, a proband with microcephaly was identified to have a homozygous missense variant in *TTI2*, further implicating ATR, HSP90 and TTT complex in microcephaly [57]. Given that PPP5C is highly expressed in the mammalian brain, and overexpression and knockdown studies showed hyper- and hypo-cellular proliferation, respectively, it is tempting to hypothesize that a reduction of PPP5C function as seen with the variant Ala47Thr could lead to microcephaly and other neurodevelopmental abnormalities observed in the proband [5, 14]. Moreover, it is also intriguing to note the parallel between the proband’s failure to thrive and the low body weight of Ppp5c^-/-^ mice [23]. However, the *Ppp5c* knockout mouse is not a perfect model of human phenotypes as it does not have reported microcephaly or intellectual disabilities. It has been suggested that mouse models of microcephaly can have limits, as the human brain is much larger when compared to the body, and mice do not have structures such as gyri and encephalization [51]. Additionally, a recent study looked at the correlation of phenotypes between known human disease gene phenotypes and knockout mouse model phenotypes, of which 49.53% do not match [58]. A recent study also demonstrated that there are differences in gene expression during the developmental timeline of humans as compared to mice and other model organisms (with approximately 15% of genes expressed in the brain during development having different expression patterns between mice and humans), which could explain differences in phenotype between human patients and mouse models [59].

Numerous chemical mutagenesis screens have been performed in *C. elegans* and other model organisms generating large collections of historical or legacy missense mutations. These missense mutations, as they disrupt protein function, provide functional information for the corresponding variant in the human ortholog or homolog. For example, the myosin heavy chain mutation *unc-54(st132)* Glu524Lys, isolated as a dominant extragenic suppressor of the *unc-22* (human titan ortholog) twitching phenotype [60, 61] is at the same position and the same change in the myosin motor domain as *MYH7* Glu525Lys, a pathogenic variant in dilated cardiomyopathy [62]. This *unc-54* legacy mutation provides experimental support for the *MYH7* variant being damaging, where the data were available more than 10 years before identification of the variant from nextgeneration sequencing of a patient cohort.

We have examined a collection of 25 *pph-5* legacy missense mutations identified from forward genetic screens [29, 30]. For 10 of these the residue is identical or a conservative substitution in *PPP5C,* there are no instances of variants at these residues in gnomAD, and all the *pph-5* mutations are strong hypomorphs or complete loss of function, and thus likely to contribute to human phenotype if observed in *PPP5C,* under a model of dominant presentation due to haploinsufficiency (Table 1). The 10 variants have yet to be reported in the human population in ClinVar (search date 12/27/2021). These 10 legacy mutations provide functional data that the missense changes are damaging in *C. elegans* and, by extension, when observed in humans likely to contribute to disease. Given the potential utility of model organism legacy missense mutation findings to human gene-variant analysis, model organism databases (WormBase, FlyBase, Alliance of Genome Resources, etc.) should aggregate and display such data similar to that shown in Table 1 and make that information available to databases such as ClinVar and MARRVEL.

In summary, *PPP5C* p.Ala47Thr, a *de novo* heterozygous missense gene-variant identified in a UDN proband, was modeled in *C. elegans. In vivo* functional analyses indicated that the *pph-5* Ala48Thr variant is damaging, similar in extent to the complete *pph-5* loss of function. By extension, our data suggest that *PPP5C* p.Ala47Thr is also damaging, likely resulting in a strong hypomorph or complete loss of function. Given that PPP5C, HSP90 and client proteins are important for the development of the brain, it is likely that the *PPP5C* p.Ala47Thr loss-of-function variant is responsible for one or more of the proband’s phenotypes, such as microcephaly and/or the other neurodevelopmental disorders.

## ACKNOWLEDGEMENTS

Research reported in this manuscript was supported by the NIH Common Fund, through the Office of Strategic Coordination/Office of the NIH Director under Award Number U54 NS108251 (TS and Lila Solnica-Krezel) and U01HG007709. Funding was also provided by the National Institutes of Health (R01 GM11447, JNB), the Children’s Discovery Institute, St Louis Children’s Hospital Foundation (GAS and SCP). Some strains were provided by the CGC, which is funded by the NIH Office of Research Infrastructure Programs (P40 OD010440). The content of this manuscript is solely the responsibility of the authors and does not necessarily represent the official views of the National Institutes of Health.

## Author Contributions

S.M.F., T.S., and S.C.P. designed research; S.M.F, J.N.B., and H.H. performed research; S.M.F., J.A.R., L.C.B., L.E., S.L., R.A., J.N.B., D.B., G.A.S., T.S., and S.C.P. analyzed data; and S.M.F, T.S., and S.C.P. wrote the paper.

## CONSORTIA (Members of the Undiagnosed Diseases Network)

Maria T. Acosta, Margaret Adam, David R. Adams, Justin Alvey, Laura Amendola, Ashley Andrews, Euan A. Ashley, Mahshid S. Azamian, Carlos A. Bacino, Guney Bademci, Ashok Balasubramanyam, Dustin Baldridge, Jim Bale, Michael Bamshad, Deborah Barbouth, Pinar Bayrak-Toydemir, Anita Beck, Alan H. Beggs, Edward Behrens, Gill Bejerano, Hugo Bellen, Jimmy Bennet, Beverly Berg-Rood, Jonathan A. Bernstein, Gerard T. Berry, Anna Bican, Stephanie Bivona, Elizabeth Blue, John Bohnsack, Devon Bonner, Lorenzo Botto, Brenna Boyd, Lauren C. Briere, Elly Brokamp, Gabrielle Brown, Elizabeth A. Burke, Lindsay C. Burrage, Manish J. Butte, Peter Byers, William E. Byrd, John Carey, Olveen Carrasquillo, Thomas Cassini, Ta Chen Peter Chang, Sirisak Chanprasert, Hsiao-Tuan Chao, Gary D. Clark, Terra R. Coakley, Laurel A. Cobban, Joy D. Cogan, Matthew Coggins, F. Sessions Cole, Heather A. Colley, Cynthia M. Cooper, Heidi Cope, William J. Craigen, Andrew B. Crouse, Michael Cunningham, Precilla D’Souza, Hongzheng Dai, Surendra Dasari, Joie Davis, Jyoti G. Dayal, Matthew Deardorff, Esteban C. Dell’Angelica, Katrina Dipple, Daniel Doherty, Naghmeh Dorrani, Argenia L. Doss, Emilie D. Douine, Laura Duncan, Dawn Earl, David J. Eckstein, Lisa T. Emrick, Christine M. Eng, Cecilia Esteves, Marni Falk, Liliana Fernandez, Elizabeth L. Fieg, Laurie C. Findley, Paul G. Fisher, Brent L. Fogel, Irman Forghani, William A. Gahl, Ian Glass, Bernadette Gochuico, Rena A. Godfrey, Katie Golden-Grant, Madison P. Goldrich, Alana Grajewski, Irma Gutierrez, Don Hadley, Sihoun Hahn, Rizwan Hamid, Kelly Hassey, Nichole Hayes, Frances High, Anne Hing, Fuki M. Hisama, Ingrid A. Holm, Jason Hom, Martha Horike-Pyne, Alden Huang, Yong Huang, Wendy Introne, Rosario Isasi, Kosuke Izumi, Fariha Jamal, Gail P. Jarvik, Jeffrey Jarvik, Suman Jayadev, Orpa Jean-Marie, Vaidehi Jobanputra, Lefkothea Karaviti, Jennifer Kennedy, Shamika Ketkar, Dana Kiley, Gonench Kilich, Shilpa N. Kobren, Isaac S. Kohane, Jennefer N. Kohler, Deborah Krakow, Donna M. Krasnewich, Elijah Kravets, Susan Korrick, Mary Koziura, Seema R. Lalani, Byron Lam, Christina Lam, Grace L. LaMoure, Brendan C. Lanpher, Ian R. Lanza, Kimberly LeBlanc, Brendan H. Lee, Hane Lee, Roy Levitt, Richard A. Lewis, Pengfei Liu, Xue Zhong Liu, Nicola Longo, Sandra K. Loo, Joseph Loscalzo, Richard L. Maas, Ellen F. Macnamara, Calum A. MacRae, Valerie V. Maduro, Rachel Mahoney, Bryan C. Mak, May Christine V. Malicdan, Laura A. Mamounas, Teri A. Manolio, Rong Mao, Kenneth Maravilla, Ronit Marom, Gabor Marth, Beth A. Martin, Martin G. Martin, Julian A. Martínez-Agosto, Shruti Marwaha, Jacob McCauley, Allyn McConkie-Rosell, Alexa T. McCray, Elisabeth McGee, Heather Mefford, J. Lawrence Merritt, Matthew Might, Ghayda Mirzaa, Eva Morava, Paolo M. Moretti, John J. Mulvihill, Mariko Nakano-Okuno, Stan F. Nelson, John H. Newman, Sarah K. Nicholas, Deborah Nickerson, Shirley Nieves-Rodriguez, Donna Novacic, Devin Oglesbee, James P. Orengo, Laura Pace, Stephen C. Pak, J. Carl Pallais, Christina GS. Palmer, Jeanette C. Papp, Neil H. Parker, John A. Phillips III, Jennifer E. Posey, Lorraine Potocki, Barbara N. Pusey, Aaron Quinlan, Wendy Raskind, Archana N. Raja, Deepak A. Rao, Anna Raper, Genecee Renteria, Chloe M. Reuter, Lynette Rives, Amy K. Robertson, Lance H. Rodan, Jill A. Rosenfeld, Natalie Rosenwasser, Francis Rossignol, Maura Ruzhnikov, Ralph Sacco, Jacinda B. Sampson, Mario Saporta, C. Ron Scott, Judy Schaechter, Timothy Schedl, Kelly Schoch, Daryl A. Scott, Vandana Shashi, Jimann Shin, Rebecca Signer, Edwin K. Silverman, Janet S. Sinsheimer, Kathy Sisco, Edward C. Smith, Kevin S. Smith, Emily Solem, Lilianna Solnica-Krezel, Ben Solomon, Rebecca C. Spillmann, Joan M. Stoler, Jennifer A. Sullivan, Kathleen Sullivan, Angela Sun, Shirley Sutton, David A. Sweetser, Virginia Sybert, Holly K. Tabor, Amelia L. M. Tan, Queenie K.-G. Tan, Mustafa Tekin, Fred Telischi, Willa Thorson, Cynthia J. Tifft, Camilo Toro, Alyssa A. Tran, Brianna M. Tucker, Tiina K. Urv, Adeline Vanderver, Matt Velinder, Dave Viskochil, Tiphanie P. Vogel, Colleen E. Wahl, Stephanie Wallace, Nicole M. Walley, Melissa Walker, Jennifer Wambach, Jijun Wan, Lee-kai Wang, Michael F. Wangler, Patricia A. Ward, Daniel Wegner, Monika Weisz-Hubshman, Mark Wener, Tara Wenger, Katherine Wesseling Perry, Monte Westerfield, Matthew T. Wheeler, Jordan Whitlock, Lynne A. Wolfe, Jeremy D. Woods, Kim Worley, Changrui Xiao, Shinya Yamamoto, John Yang, Diane B. Zastrow, Zhe Zhang, Chunli Zhao, Stephan Zuchner

## REFERENCES

1. Ali, A., et al., Requirement of protein phosphatase 5 in DNA-damage-induced ATM activation. Genes & development, 2004. 18(3): p. 249–254.

2. Kaziales, A., et al., Glucocorticoid receptor complexes form cooperatively with the Hsp90 co-chaperones Pp5 and FKBPs. Scientific reports, 2020. 10(1): p. 1–16.

3. Yong, W., et al., Mice lacking protein phosphatase 5 are defective in ataxia telangiectasia mutated (ATM)-mediated cell cycle arrest. Journal of Biological Chemistry, 2007. 282(20): p. 14690–14694.

4. Zhang, J., et al., Protein phosphatase 5 is required for ATR-mediated checkpoint activation. Molecular and cellular biology, 2005. 25(22): p. 9910–9919.

5. Zuo, Z., N.M. Dean, and R.E. Honkanen, Serine/threonine protein phosphatase type 5 acts upstream of p53 to regulate the induction of p21WAF1/Cip1 and mediate growth arrest. Journal of Biological Chemistry, 1998. 273(20): p. 12250–12258.

6. Wechsler, T., et al., DNA-PKcs function regulated specifically by protein phosphatase 5. Proceedings of the National Academy of Sciences, 2004. 101(5): p. 1247–1252.

7. Partch, C.L., et al., Posttranslational regulation of the mammalian circadian clock by cryptochrome and protein phosphatase 5. Proceedings of the National Academy of Sciences, 2006. 103(27): p. 10467–10472.

8. Liu, F., et al., Dephosphorylation of tau by protein phosphatase 5: impairment in Alzheimer’s disease. Journal of Biological Chemistry, 2005. 280(3): p. 1790–1796.

9. Hinds Jr, T.D. and E.R. Sánchez, Protein phosphatase 5. The international journal of biochemistry & cell biology, 2008. 40(11): p. 2358–2362.

10. Bahl, R., et al., Localization of protein Ser/Thr phosphatase 5 in rat brain. Molecular brain research, 2001. 90(2): p. 101–109.

11. Becker, W., et al., Distribution of the mRNA for protein phosphatase T in rat brain. Molecular brain research, 1996. 36(1): p. 23–28.

12. Becker, W., et al., Molecular cloning of a protein serine/threonine phosphatase containing a putative regulatory tetratricopeptide repeat domain. Journal of Biological Chemistry, 1994. 269(36): p. 22586–22592.

13. Chinkers, M., Targeting of a distinctive protein-serine phosphatase to the protein kinase-like domain of the atrial natriuretic peptide receptor. Proceedings of the National Academy of Sciences, 1994. 91(23): p. 11075–11079.

14. Lv, J.-M., et al., PPP5C promotes cell proliferation and survival in human prostate cancer by regulating of the JNK and ERK1/2 phosphorylation. OncoTargets and therapy, 2018. 11: p. 5797.

15. Chen, M., et al., Disruption of serine/threonine protein phosphatase 5 inhibits tumorigenesis of urinary bladder cancer cells. International journal of oncology, 2017. 51(1): p. 39–48.

16. Urban, G., et al., Identification of an estrogen-inducible phosphatase (PP5) that converts MCF-7 human breast carcinoma cells into an estrogen-independent phenotype when expressed constitutively. Journal of Biological Chemistry, 2001. 276(29): p. 27638–27646.

17. Xie, J., et al., PP5 (PPP5C) is a phosphatase of Dvl2. Scientific reports, 2018. 8(1): p. 1–15.

18. Barnes, P.J., Anti-inflammatory actions of glucocorticoids: molecular mechanisms. Clinical science, 1998. 94(6): p. 557–572.

19. Sapolsky, R.M., L.M. Romero, and A.U. Munck, How do glucocorticoids influence stress responses? Integrating permissive, suppressive, stimulatory, and preparative actions. Endocrine reviews, 2000. 21(1): p. 55–89.

20. Sugimoto, K., Branching the Tel2 pathway for exact fit on phosphatidylinositol 3-kinase-related kinases. Current genetics, 2018. 64(5): p. 965–970.

21. Abraham, R.T., PI 3-kinase related kinases:’big’players in stress-induced signaling pathways. DNA repair, 2004. 3(8-9): p. 883–887.

22. Amable, L., et al., Disruption of serine/threonine protein phosphatase 5 (PP5: PPP5c) in mice reveals a novel role for PP5 in the regulation of ultraviolet light-induced phosphorylation of serine/threonine protein kinase Chk1 (CHEK1). Journal of Biological Chemistry, 2011. 286(47): p. 40413–40422.

23. Grankvist, N., et al., Serine/threonine protein phosphatase 5 regulates glucose homeostasis in vivo and apoptosis signalling in mouse pancreatic islets and clonal MIN6 cells. Diabetologia, 2012. 55(7): p. 2005–2015.

24. Grankvist, N., et al., Genetic disruption of protein phosphatase 5 in mice prevents high-fat diet feeding-induced weight gain. FEBS letters, 2013. 587(23): p. 3869–3874.

25. Brown, L., E.B. Borthwick, and P.T. Cohen, Drosophila protein phosphatase 5 is encoded by a single gene that is most highly expressed during embryonic development. Biochimica et Biophysica Acta (BBA)-Gene Structure and Expression, 2000. 1492(2-3): p. 470–476.

26. Chen, F., et al., Multiple protein phosphatases are required for mitosis in Drosophila. Current Biology, 2007. 17(4): p. 293–303.

27. Jeong, J.-Y., et al., Characterization of Saccharomyces cerevisiae protein Ser/Thr phosphatase T1 and comparison to its mammalian homolog PP5. BMC Cell Biology, 2003. 4(1): p. 1–13.

28. Zheng, C., et al., Opposing effects of an F-box protein and the HSP90 chaperone network on microtubule stability and neurite growth in Caenorhabditis elegans. Development, 2020. 147(12): p. dev189886.

29. Richie, C.T., et al., Protein phosphatase 5 is a negative regulator of separase function during cortical granule exocytosis in C. elegans. Journal of Cell Science, 2011. 124(17): p. 2903–2913.

30. Melesse, M., et al., Genetic identification of separase regulators in Caenorhabditis elegans. G3: Genes, Genomes, Genetics, 2018. 8(2): p. 695–705.

31. Karczewski, K.J., et al., The mutational constraint spectrum quantified from variation in 141,456 humans. Nature, 2020. 581(7809): p. 434–443.

32. Huang, H., et al., A dominant negative variant of RAB5B disrupts maturation of surfactant protein B and surfactant protein C. Proceedings of the National Academy of Sciences, 2022. 119(6).

33. Boulin, T., et al., Functional analysis of a de novo variant in the neurodevelopment and generalized epilepsy disease gene NBEA. Molecular Genetics and Metabolism, 2021. 134(1-2): p. 195–202.

34. Suzuki, H., et al., Biallelic loss of OTUD7A causes severe muscular hypotonia, intellectual disability, and seizures. Am J Med Genet A, 2021. 185(4): p. 1182–1186.

35. Brenner, S., The genetics of Caenorhabditis elegans. Genetics, 1974. 77(1): p. 71–94.

36. Satterstrom, F.K., et al., Large-scale exome sequencing study implicates both developmental and functional changes in the neurobiology of autism. Cell, 2020. 180(3): p. 568–584. e23.

37. Chen, M.-S., et al., The tetratricopeptide repeat domain of protein phosphatase 5 mediates binding to glucocorticoid receptor heterocomplexes and acts as a dominant negative mutant. Journal of Biological Chemistry, 1996. 271(50): p. 32315–32320.

38. Kang, H., et al., Identification of amino acids in the tetratricopeptide repeat and C-terminal domains of protein phosphatase 5 involved in autoinhibition and lipid activation. Biochemistry, 2001. 40(35): p. 10485–10490.

39. Karney-Grobe, S., et al., HSP90 is a chaperone for DLK and is required for axon injury signaling. Proceedings of the National Academy of Sciences, 2018. 115(42): p. E9899–E9908.

40. Vaughan, C.K., et al., Hsp90-dependent activation of protein kinases is regulated by chaperone-targeted dephosphorylation of Cdc37. Molecular cell, 2008. 31(6): p. 886–895.

41. Haslbeck, V., et al., The activity of protein phosphatase 5 towards native clients is modulated by the middle-and C-terminal domains of Hsp90. Scientific reports, 2015. 5(1): p. 1–16.

42. Rand, J., Acetylcholine (January 30, 2007), WormBook, ed. The C. elegans Research Community, WormBook, doi/10.1895/wormbook.1.131.1. 2007.

43. Richmond, J.E. and E.M. Jorgensen, One GABA and two acetylcholine receptors function at the C. elegans neuromuscular junction. Nature neuroscience, 1999. 2(9): p. 791–797.

44. Sun, Y., et al., The F-box protein MEC-15 (FBXW9) promotes synaptic transmission in GABAergic motor neurons in C. elegans. PloS one, 2013. 8(3): p. e59132.

45. Loria, P.M., J. Hodgkin, and O. Hobert, A conserved postsynaptic transmembrane protein affecting neuromuscular signaling in Caenorhabditis elegans. Journal of Neuroscience, 2004. 24(9): p. 2191–2201.

46. Siomos, M.F., et al., Separase is required for chromosome segregation during meiosis I in Caenorhabditis elegans. Current Biology, 2001. 11(23): p. 1825–1835.

47. Bembenek, J.N., et al., Cortical granule exocytosis in C. elegans is regulated by cell cycle components including separase. 2007.

48. Schopf, F.H., M.M. Biebl, and J. Buchner, The HSP90 chaperone machinery. Nature reviews Molecular cell biology, 2017. 18(6): p. 345–360.

49. Takai, H., et al., Tel2 structure and function in the Hsp90-dependent maturation of mTOR and ATR complexes. Genes & development, 2010. 24(18): p. 2019–2030.

50. O’Driscoll, M., A. Jackson, and P.A. Jeggo, Microcephalin: a causal link between impaired damage response signalling and microcephaly. Cell cycle, 2006. 5(20): p. 2339–2344.

51. O’Driscoll, M. and P.A. Jeggo, The role of the DNA damage response pathways in brain development and microcephaly: insight from human disorders. DNA repair, 2008. 7(7): p. 1039–1050.

52. Abner, C.W. and P.J. McKinnon, The DNA double-strand break response in the nervous system. DNA repair, 2004. 3(8-9): p. 1141–1147.

53. Orii, K.E., et al., Selective utilization of nonhomologous end-joining and homologous recombination DNA repair pathways during nervous system development. Proceedings of the National Academy of Sciences, 2006. 103(26): p. 10017–10022.

54. Takai, H., et al., Tel2 regulates the stability of PI3K-related protein kinases. Cell, 2007. 131(7): p. 1248–1259.

55. Hurov, K.E., C. Cotta-Ramusino, and S.J. Elledge, A genetic screen identifies the Triple T complex required for DNA damage signaling and ATM and ATR stability. Genes & development, 2010. 24(17): p. 1939–1950.

56. You, J., et al., A syndromic intellectual disability disorder caused by variants in TELO2, a gene encoding a component of the TTT complex. The American Journal of Human Genetics, 2016. 98(5): p. 909–918.

57. Picher-Martel, V., et al., Whole-exome sequencing identifies homozygous mutation in TTI2 in a child with primary microcephaly: a case report. BMC neurology, 2020. 20(1): p. 1–6.

58. Cacheiro, P., M.A. Haendel, and D. Smedley, New models for human disease from the International Mouse Phenotyping Consortium. Mammalian Genome, 2019. 30(5): p. 143–150.

59. Cardoso-Moreira, M., et al., Developmental gene expression differences between humans and mammalian models. Cell reports, 2020. 33(4): p. 108308.

60. Moerman, D.G., et al., Mutations in the unc-54 myosin heavy chain gene of Caenorhabditis elegans that alter contractility but not muscle structure. Cell, 1982. 29(3): p. 773–781.

61. Moerman, D., A. Fire, and D. In Riddle, C. elegans II. 1997, Cold Spring Harbor Laboratory Press, New York.

62. Lakdawala, N.K., et al., Genetic testing for dilated cardiomyopathy in clinical practice. Journal of cardiac failure, 2012. 18(4): p. 296–303.

