## Supplemental information for "Functional analysis of a novel *de novo* variant in *PPP5C* associated with microcephaly, seizures, and developmental delay"

### **SUPPORTING INFORMATION**

**Supplementary Figure S1**

**Supplementary Table S1**

**Supplementary Table S2**

**Supplementary Table S3**

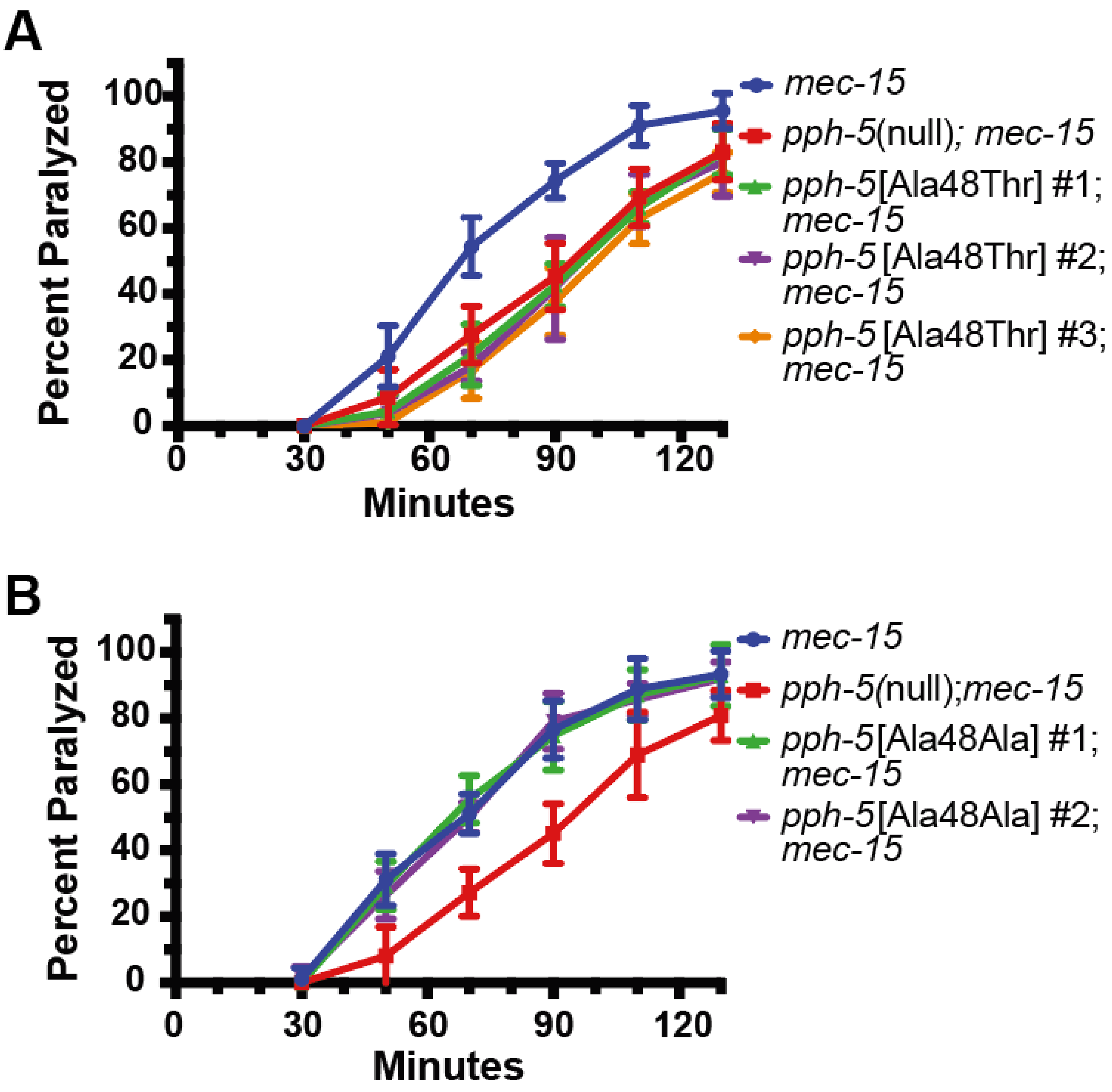


Figure S1. Independent CRISPR-edited strains behave similarly. (A) Quantification of percent animals paralyzed after exposure to aldicarb of all three patient variant lines *pph-5*[Ala48Thr]; *mec-15* compared to *mec-15* and to *pph-5(null); mec-15*. All three patient variant lines show the same rate of paralysis as each other and as *pph-5(null); mec-15*. (B) Quantification of percent animals paralyzed after exposure to aldicarb of both control edit lines (Ala48Ala); *mec-15* compared to *mec-15* and to *pph-5(null);mec-15*. Both control edit lines with *mec-15* paralyze at the same rate as each other and *mec-15.*

**Table S1. Variants identified by research analysis of the trio exome sequencing**

| Gene Name | Genomic Coordinates (Hg19) | Transcript | Variant | gnomAD | Inheritance | Notes |
|---|---|---|---|---|---|---|
| *FSIP2* | 2:186678720C>T | NM_173651.2 | c.20543C>T ( p.Thr6848Met) | 151/276352 | Inherited from father | CADD<1 |
| *FSIP2* | 2:186654475A>G | NM_173651.2 | c.2879A>G (p.Asn960Ser) | 24/167422 | Inherited from mother | CADD=5.7 |
| *MICAL3* | 22:18304892G>A | NM_015241.2 | c.3352C>T (p.Arg1118Cys) | 9/280386 | Inherited from father | CADD=16.9 |
| *MICAL3* | 22:18347692C>T | NM_015241.2 | c.2578G>A (p.Val860Met) | 194/278050 | Inherited from mother | CADD=3.2 |
| *SPTB* | 14:65253409A>G | NM_001024858.2 | c.3274T>C (p.Ser1092Pro) | Not present | Inherited from mother[1] | CADD=24.6 |
| *SPTB* | 14:65268075G>A | NM_000347.5 | c.691C>T (p.Arg231Trp) | 22/282838 | Inherited from father | CADD=24.5 |
| *PPP5C* | 19:46857022G>A | NM_001204284.1 | c.139G>A (p.Ala47Thr) | Not present | De Novo | CADD=29.4 |

*De novo* variants that are not observed in gnomAD and biallelic variants in the coding region or near intron-exon boundaries with allele frequency of less than 1% (and not homozygous in gnomAD) are listed above. [1]Father has one read with the variant.

**Table S2. List of *C. elegans* strains used in this study**

| Strain name | genotype | Description |
|---|---|---|
| RB2522 | *pph-5(ok3498)* | Control for *pph-5* loss of function |
| TU5237 | *mec-15(u1042) II; uIs115* | Control line used for aldicarb assay and PLM-AN growth measurement |
| | *uIs115* | Crossed TU5237 with VC2010 to get uIS115 on its own as a WT control for PLM-AN growth measurement |
| UDN100160 | *pph-5(udn90) V; mec-15(u1042) II; uIs115* | Ala48Thr proband variant line #1. Used for aldicarb assay and PLM-AN growth measurement |
| UDN100161 | *pph-5(udn91) V; mec-15(u1042) II; uIs115* | Ala48Thr proband variant line #2. Used for aldicarb assay and PLM-AN growth measurement |
| UDN100162 | *pph-5(udn92) V; mec-15(u1042) II; uIs115* | Ala48Thr proband variant line #3- excluded from experiments as not statistically different from other two proband variant lines |
| UDN100163 | *pph-5(udn93) V* | Ala48Ala control line #1. Used for aldicarb assay and PLM-AN growth measurement |
| UDN100164 | *pph-5(udn94) V* | Ala48Ala control line #2- excluded from experiments as not statistically different from the other control edit line or VC2010 |
| | *sep-1(e2406) I* | |
| | *sep-1(e2406) I; pph-5(udn90) V* | Ala48Thr proband variant line #1. Used for embryonic lethality experiment |
| | *sep-1(e2406) I; pph-5(udn91) V* | Ala48Ala proband variant line #2. Used for embryonic lethality experiment |
| | *sep-1(e2406) I; pph-5(udn93) V* | Ala48Ala control line #1. Used for embryonic lethality experiment |
| | *sep-1(e2406) I; pph-5(udn94) V* | Ala48Ala control line #2. Used for embryonic lethality experiment |

**Table S3. Single-stranded DNA repair template sequences.**

| oligo | Single-stranded DNA repair template sequence |
|---|---|
| Proband variant edit repair template | TTGATGAATTTTAAAGTATTTTCCTGCTGCAGATCAAGTGTAtGAtGTa**aCC**GCgGACCTCTACTCTGTCGCAATTGAGATTCATCCGACGGCGGTTCT |
| Control edit repair template | TTTGATGAATTTTAAAGTATTTTCCTGCTGCAGATCAAGTGTAtGAtGTa**GCC**GCgGACCTCTACTCTGTCGCAATTGAGATTCATCCGACGGCGGTTCT |

Lower case letters indicate changed nucleotides from the reference sequence. AvaII restriction site added. PAM sequence underlined. Ala48Ala/Thr edits in bold
